# SPOT: a web-tool enabling Swift Profiling Of Transcriptomes

**DOI:** 10.1101/2021.03.03.433767

**Authors:** Elias Farr, Julia M. Sattler, Friedrich Frischknecht

## Abstract

The increasing number of single cell and bulk RNAseq data sets describing complex gene expression profiles in different organisms, organs or cell types calls for an intuitive tool allowing rapid comparative analysis. Here we present Swift Profiling Of Transcriptomes (SPOT) as a web tool that allows not only differential expression analysis but also fast ranking of genes fitting transcription profiles of interest. Based on a heuristic approach the spot algorithm ranks the genes according to their proximity to the user-defined gene expression profile of interest. The best hits are visualized as a table, bar chart or dot plot and can be exported as an Excel file. While the tool is generally applicable, we tested it on RNAseq data from malaria parasites that undergo multiple stage transformations during their complex life cycle as well as on data from multiple human organs during development and cell lines infected by the SARS-CoV-2 virus. SPOT should enable non-bioinformaticians to easily analyse their own and any available dataset.

## 1 Introduction

To understand cell and organ-specific functionality, gene expression patterns are often determined as a first step before specific genes are investigated in more detail. Many researchers are interested to identify genes that are specifically expressed in certain entities such as cell types, developmental stages, tissues or species where their encoded proteins can play diverse or similar roles. Different tools exist to visualize profiles of known genes or to perform differential expression analysis (DEA) (Ge et al. 2018; Papatheodorou et al. 2020; Reyes et al. 2019).

However, there is no intuitive tool to extract and visualize gene expression patterns matching user-defined gene expression profiles of interest, POI (Figure 1A). SPOT addresses this need (Figure 1B, C): sliders on the application’s control panel allow users to enter their own POI across the desired entities of a data set. This slider-entity connection is the basis for a ranking process that is based on the proximity to the user-defined gene expression POI, which is updated every time a slider is moved. The results of the spot ranking algorithm are visualized in the main panel as table, bar chart or dot blot (Figure 1C).

**Figure 1.**
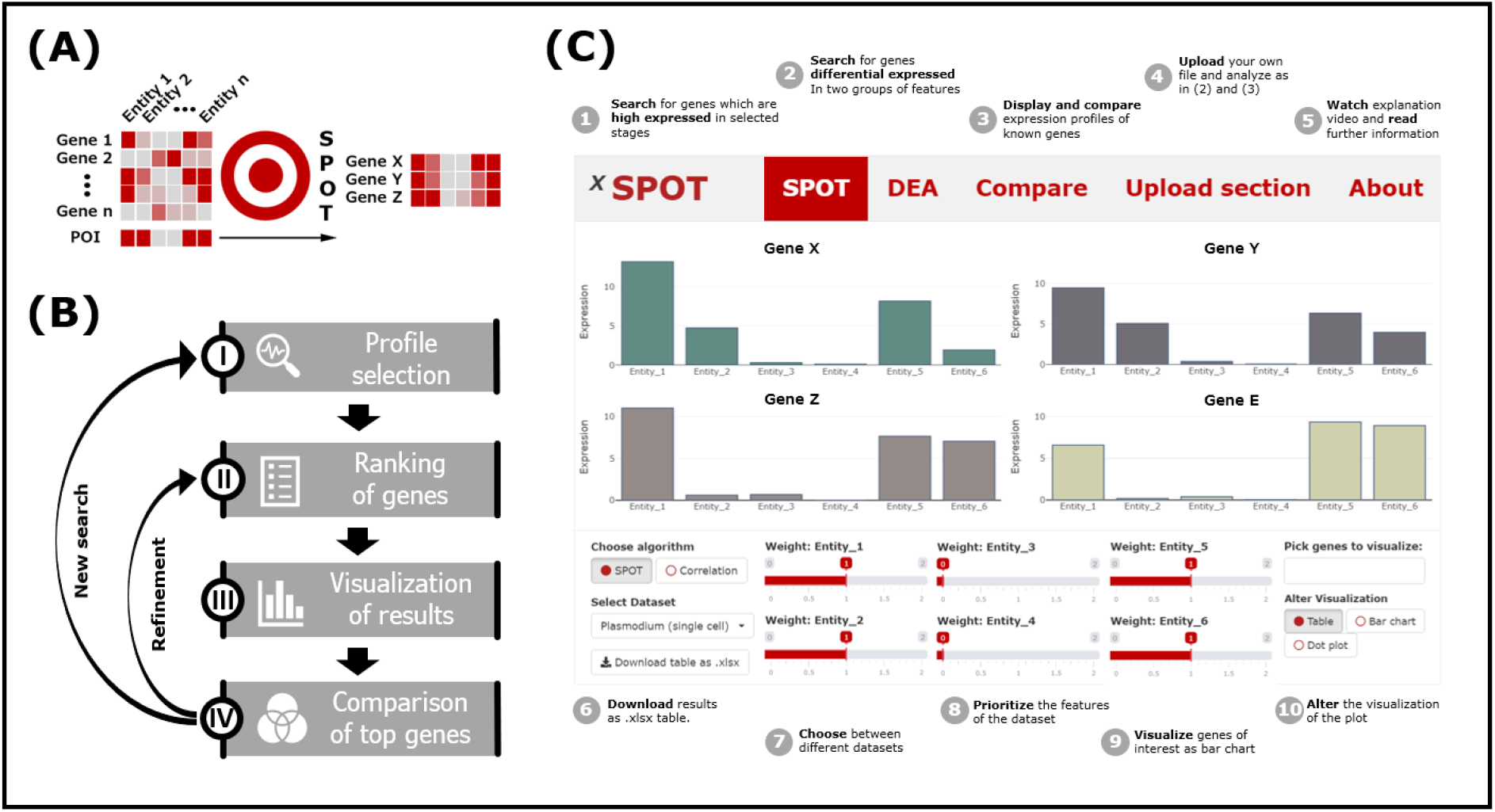
(A) Logic and (B) workflow of SPOT. (C) Cartoon version of the SPOT web interface highlighting specific functions

For the proof-of-concept we pre-installed data in SPOT from studies on organ development of Homo sapiens (bulkRNAseq) (Cardoso-Moreira et al. 2019) and developmental stages of the rodent infecting malaria parasite Plasmodium berghei (scRNAseq) (Howick et al. 2019) as well as SARS-CoV-2 infected human cell lines (Wyler et al. 2021). The general workflow and the interface design can be seen in Figure 1, while example predictions and comparison with state-of-the-art methods are available in the supplemental material.

## 2 Methods and features

SPOT is a web-based R shiny app, which mainly consists of a visualization and a control panel. While the user can make selections in widgets such as tables, sliders, drop-down menus or buttons in the control panel, the visualization panel depicts the results of the given input. For each analysis variant, the app has a section with its own functionalities in the lower panel and visualization methods in the upper panel.

### 2.1 SPOT Expression Profile

In this proof-of-concept version of SPOT expression profiles from the three implemented sets of data can be extracted. To this end, the POI of each entity (e.g. cell type, stage of development, organ, infected vs non-infected cell) can be defined via a slider with values from 0 to 2. spot ranks the genes according to the match of their expression to the POI as reflected by the slider constellation. The top ranked genes can be displayed as table, bar chart or dot plot. In case of a single cell dataset, the dot plot visualizes the average of cells in which gene expression is detected as dot circumference – and the average expression as dot colour. For bulk RNAseq data, gene expression is visualized as circumference of a dot. For further analysis, the results can be downloaded as Excel file. Also links to the database entries of the individual genes are provided. A detailed definition of the spot algorithm is available in the supplementary information.

### 2.2 DEA

For users interested in a more precise but less interactive candidate search, we have also implemented a differential expression analysis (DEA) tool based on Seurat, MAST and DESeq2 (Stuart et al., 2019; Finak et al. 2015; Anders and Huber, 2010). Similar to SPOT, the user can select the POI of entities in the lower panel. Due to its simplicity a Wilcoxon rank sum test between the selected and unselected entities is performed as default. As more time-consuming methods MAST (for single cells) and DESeq2 are available for ranking (Figure S1). For visualization the same charts are available as for the SPOT section.

### 2.3 Comparing profiles

Gene expression values of three pre-installed data sets are listed as a HTML table in the lower panel enabling the selection of single genes for visualization. The table can be searched with a term, filtered by multiple attributes (e.g. name, expression values) or subdivided by expression value ranges. Each selected gene is displayed as an interactive box plot in the visualization panel, which allows quick profile comparisons e.g. of already known genes with the results of the performed analysis.

### 2.4 Analysis of additional datasets

To allow users to examine their own records or published datasets, data files (.csv/.xlsx) can be readily uploaded. Input data sets should list entities in the first row and gene names in the first column. The shiny app then creates a slider for each column name (entity) in the first row. For single cell datasets the cell identities should be listed in the second row. Uploaded datasets are visualized in the same way as described under 2.1 - 2.3.

## 3 Conclusion

SPOT provides an easy-to-use web tool for ranking genes with an expression profile that matches user-defined characteristics. Moreover, it allows swift visual comparison of multiple genes as well as differential expression analysis of datasets.

