## Supplementary material for "SPOT: a web-tool enabling Swift Profiling Of Transcriptomes": Supplementary Material_bio.pdf

### Supplementary information

### Table of Figures

### 1 Algorithm description

#### 1.1 Data pre-processing

SPOT has been preinstalled with **three datasets**, one single cell RNAseq dataset describing the whole *Plasmodium* lifecycle, a bulkRNAseq dataset containing information about human organs in several developmental stages and a bulkRNAseq dataset from Sars-CoV2 virus infected cell lines (Cardoso-Moreira, et al., 2019; Howick, et al., 2019).

For spot analysis of the **malaria cell cycle**, TMMlog normalized single cells were assigned by ShortenedLifestage4 (Howick, et al., 2019) and averaged to obtain a single expression value for every developmental stage.

**Human organ** TPM values were also condensed by averaging between multiple replicates. Since the similarity of prenatal developmental stages is high, only 3 prominent stages (4, 10, 20 weeks post conception) were kept for further analysis.

TPM values of **Sars-CoV2 infected cell lines** were calculated by the gene length specified by the publication (Wyler, et al., 2021). Samples were again condensed by averaging through the datasets for specific timepoints.

For spot and correlation ranking, data is **scaled by standardization** to obtain comparability between datasets. To lower the influence of outliers every standardized value higher than 4 was set to 4 and every standardized value below -1 was set to -1.

In contrast to *spot*, for **differential expression analysis** raw counts were used and filtered for cells having expression in more than 3 features.

#### 1.2 SPOT

The *spot* algorithm consists of two factors: (i) the difference between the weighted mean of selected (slider values > 0) or unselected entities (slider values = 0) and (ii) the difference between 1 and the mean of unselected entities. For an optimal result, both factors should be as high as possible.

The first factor measures the distance between the selected and unselected entities, by calculating the difference of their mean expression values. To enhance the influence of single entities to the difference, entities can be assigned multiple times to the regarding average. This is done by adjusting the slider values and can be useful if one selected variable has constant higher expression values than other ones. The higher the slider value, the more often the variable is assigned in the average, the lower slider value, the less often the variable is assigned.

The second factor measures how close the expression values of the unselected entities are to zero (desired in this approach). The closer they are, the less gets subtracted from one and therefore the second factor increases. Due to the scaling it is also possible to obtain negative values, for the mean.

Taken together both factors result in the *spot* score which is therefore a measure for the proximity to the user defined input. *spot* scores lie in the range of 0 to 10, the highest values obtained in these datasets are around 8.

Mathematically *spot* is defined as:

$$I := \{1, \dots, N\}$$

$$I_s := \{1 \leq i \leq N \mid column\ i\ selected\}$$

$$n_s = \# I_s$$

$$\overline{sc} = \frac{1}{n_s} \sum_{i \in I_s} \alpha_i c_i$$

$$\overline{uc} = \frac{1}{n_s} \sum_{i \in I \setminus I_s} \alpha_i c_i$$

$$spot = (\overline{sc} - \overline{uc})(1 - \overline{uc})$$

#### 1.3 Correlation

To enable subcategories between high and low expression, we implemented an option in the user interface simply calculating the Pearson correlation between the user defined profile and the genes in the dataset. As a result of the cutoff (see chapter 1.1) the highest values in the data sets are equal to 4, while the lowest generally lie close to 0. Since the sliders range from 0 to 2, the slider values are multiplied by 2 and match therefore the minima and maxima of the standardized values. This way, more complex profiles can be created with the sliders. Medium values in some entities can also be achieved (see example below). However, mean expression values in unselected columns are higher compared to *spot* or DEA approaches (Figure S2).

**Example:**

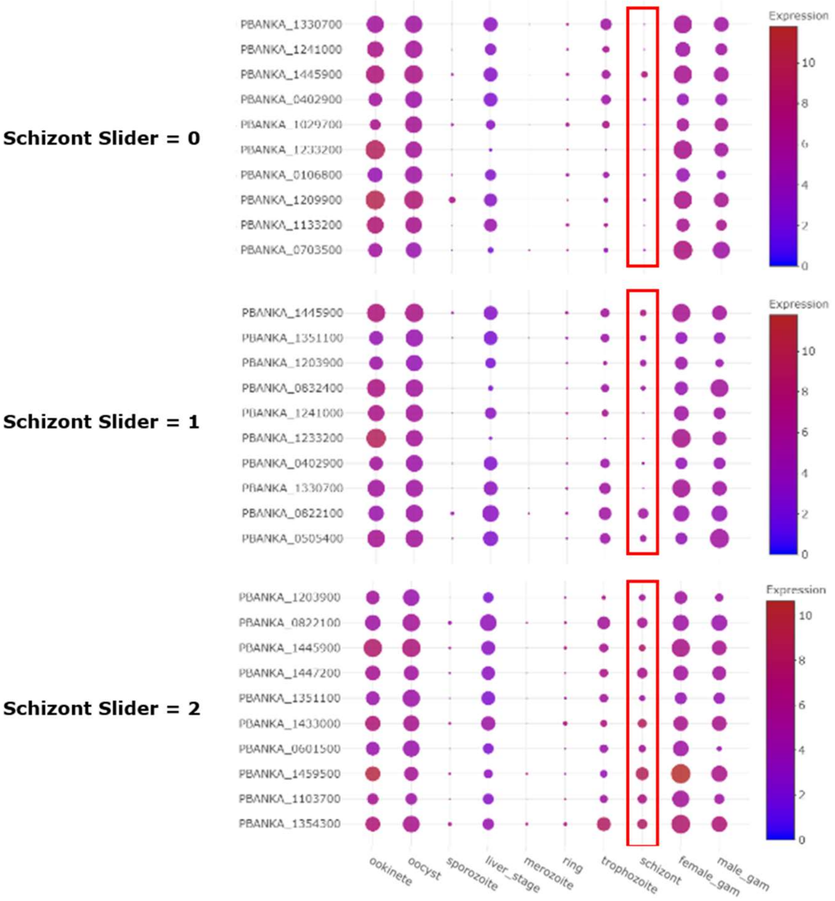

*Plasmodium* parasite expression across the complex life cycle in shown here for 10 different stages. The schizont stage describes the stage where new progeny parasites are formed within the red blood cell and is indicated with a red rectangle. While the circle size corresponds to number of single cells in which the respective gene is detected, the heat bar on the right displays the average expression.

#### 1.4 DEA

Differential expression analysis is performed with the help of the R packages Seurat, MAST and DESeq2(Anders and Huber, 2010; Finak, et al., 2015; Stuart, et al., 2019). Single cell datasets such as the data derived from the Malaria Cell Atlas are loaded to a Seurat object and identities are specified according to the ShortenedLifestage4 clustering(Howick, et al., 2019). The Seurat function Find_Markers() then compares two groups of entities entered by the user with different test methods (see Section 2). In the output table the Bonferroni corrected p value and the log fold change are displayed. If desired by users, implementation of further analysis methods is readily feasible.

### 2 Comparisons of algorithms and tools

#### 2.1 Algorithms

Since there are several approaches to detect significantly up-regulated genes, we compared the *spot* and correlation algorithm with the state-of-the-art methods of differential analysis in speed and accuracy.

The speed was tested on 3 example predictions, which are described in the further sections. We measured the time between input of the profile and output of the results with several CPUs. The results shown in Figure S1 depict an inverse association between the simplicity of the ranking method and the speed. While the *spot* and correlation ranking finished the analysis within 5 seconds, the Wilcoxon Rank Sum test outperformed the other DEA methods such as DESeq2 and Mast.

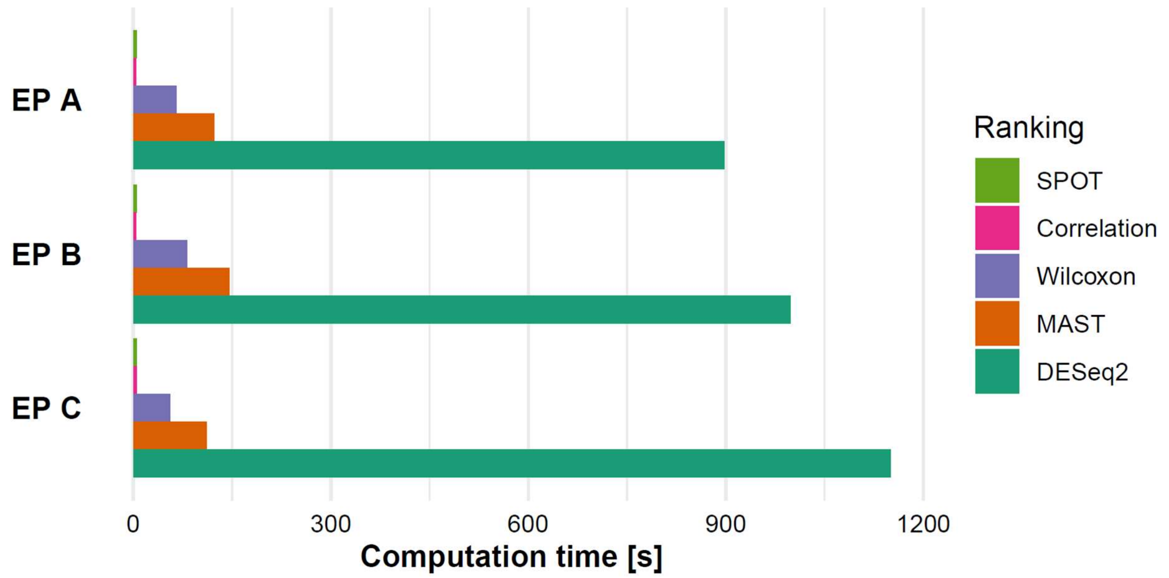

**Figure S1: *spot* algorithm outperforms DEA approaches in speed**

*spot*, Correlation and DEA methods were benchmarked for time between input in the control panel and output as table in the visualization with an AMD Ryzen 7 3700U CPU with 16 GB RAM. While *spot* and correlation stayed within 5 seconds per calculation, DEA methods ranged from 40 seconds to 19 minutes.

Accuracy tests were performed by measuring the mean expression of the best ranked candidates in selected and unselected entities of the example predictions (Figure S2). In general, the more entities are selected, the closer the values between the selected and unselected values tend to be in the example predictions. The results in the three predictions show similar accuracy of all methods with only subtle differences.

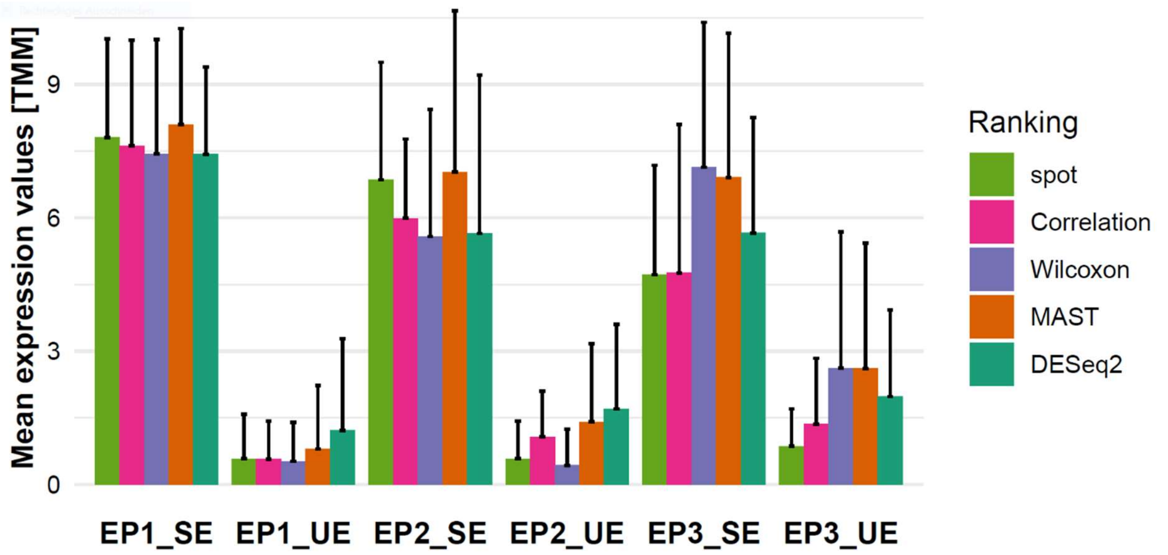

**Figure S2: Comparison of selected and unselected entities reveal best results for *spot***

Ranking accuracy in example predictions (EP) was determined through mean expression calculation of selected (SE) and unselected (UE) entities. There is overall similar accuracy revealed with MAST seemingly the best algorithm to get genes with high expression in selected entities; *spot* and the Wilcoxon test score best in unselected entities.

Since the Wilcoxon Rank Sum test is the fastest DEA methods (Figure S1), has best accuracy amongst unselected entities (Figure S2) and has also shown satisfactory results in the literature (Soneson and Robinson, 2018) it is used as the default in the web tool.

#### 2.2 Tools

Since there are multiple tools available for DEA (Ge, et al., 2018; Reyes, et al., 2019), we would like to point out the strengths and weaknesses of the approaches (Table S1). While both approaches show satisfactory results in ranking of genes (Figure S2), there are differences in speed, visualization and interface design. Although the multitude of available visualization methods gives iDEP and GENAVi an advantage over SPOT, easier handling and faster calculations make SPOT a viable alternative. SPOT might well be the better choice for users more interested in a brief overview, than in an in-depth analysis. Thus, both approaches have their advantages and disadvantages and give users the opportunity to choose according to their needs.

**Table S1: Comparison of web-tools for expression analysis**

| Parameter | DEA | SPOT |
|---|---|---|
| Accuracy | ■ Good | ■ Okay |
| Speed | ■ Bad | ■ Good |
| Visualization | ■ Good | ■ Okay |
| User interface | ■ Okay | ■ Good |

### 3 Example prediction A: *Plasmodium* liver stages

#### 3.1 Purpose

There is still no efficient malaria vaccine available (2015; Duffy and Patrick Gorres, 2020). Different approaches are currently being explored aiming to target parasites in the disease-causing blood stages or during transmission to and from the mosquito. Transmission blocking vaccines focus e.g. on generating antibodies against proteins present on the surface of gametes, which can block the transmission of *Plasmodium* to the mosquito (Carter and Chen, 1976; Gwadz, 1976), while blood stage vaccines aim e.g. at blocking parasite entry into red blood cells. In addition to such subunit vaccines also attenuated parasites are explored. Attenuated parasites are most often generated by genetic modification of genes functional during liver stage development, which can lead to a developmental arrest in hepatocytes (Kumar, et al., 2016; Mueller, et al., 2005). The specific genes are usually determined by differential expression analysis (Kaiser, et al., 2004; Matuschewski, et al., 2002). However, none of the approaches have yet succeeded in inducing sufficient protective immune responses (Duffy and Patrick Gorres, 2020). Here we present the results of a SPOT search for genes highly expressed exclusively in the liver stages. Since the genes with such expression profiles are well known (Caldelari, et al., 2019; Stanway, et al., 2019), this prediction can serve as proof-of-concept.

#### 3.2 Results

The SPOT search for genes with high expression during liver stage development revealed a top ten of genes shown in **Table S2**. Liver specific proteins 1 and 2 appear in the first 3 ranks/positions, while 3 genes with unknown functions appear in the Top 6. For all known genes except for the gametocyte specific protein and pyruvate dehydrogenase E1 component subunit beta, literature suggests strong liver stage expression (Ishino, et al., 2009; Orito, et al., 2013; Stanway, et al., 2019; Vaughan, et al., 2009). For PbANKA1003900, previously annotated as gametocyte-specific protein, recent analysis led to the suggestion of renaming it liver-specific protein due to its liver specific expression profile (Caldelari, et al., 2019; Deligianni, et al., 2018).

**Table S2: Results from *spot* ranking of liver specific genes**

| *P. Berghei* GeneID | Gene product description | *spot* score |
|---|---|---|
| PBANKA_1024600 | liver specific protein 1 | 8,01 |
| PBANKA_0519500 | conserved Plasmodium protein, unknown function | 7,75 |
| PBANKA_1003000 | liver specific protein 2 | 7,19 |
| PBANKA_1003900 | gametocyte-specific protein | 6,63 |
| PBANKA_0518900 | conserved Plasmodium membrane protein, unknown function | 5,80 |
| PBANKA_1462600 | conserved Plasmodium protein, unknown function | 3,83 |
| PBANKA_0505000 | dihydrolipoamide acyltransferase, putative | 3,62 |
| PBANKA_1310100 | pyruvate dehydrogenase E1 component subunit beta, putative | 3,58 |
| PBANKA_1125100 | 3-oxoacyl-acyl-carrier protein synthase, putative | 3,08 |
| PBANKA_1303600 | leucine carboxyl methyltransferase, putative | 3,08 |

#### 3.3 Comparison to DEA

To check the quality of the *spot* ranking, we compared the best performing genes of the *spot* ranking with those of the DEA methods. While the first 100 genes in the *spot* ranking are significantly upregulated according to the Wilcoxon test, 14 of the top 20 overlap in both rankings (Figure S3).

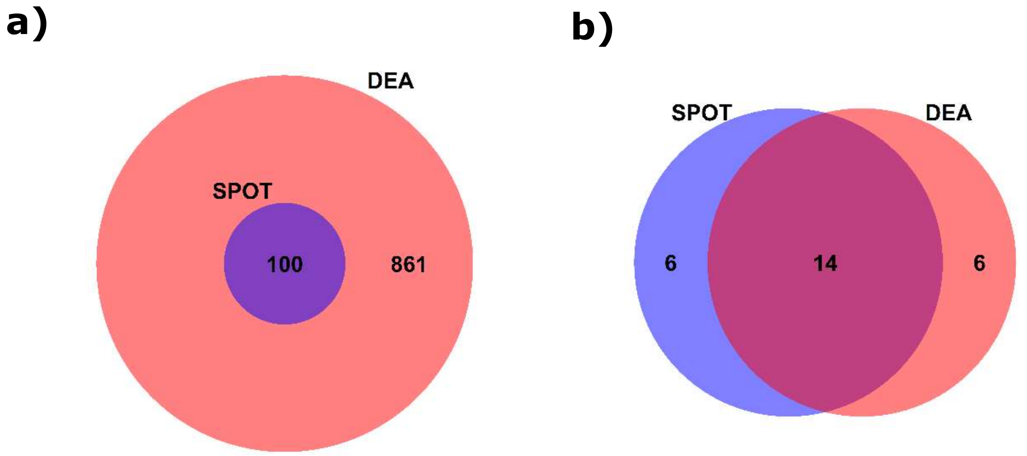

**Figure S3: First hundred genes ranked in SPOT liver stage ranking are differentially expressed**

**a)** 861 *P. berghei* genes are upregulated exclusively in the liver (p value < 0,001). The first 100 genes of the *spot* ranking overlap with these results. **b)** Two thirds (14) of the top 20 genes of both tests overlap.

The best performing candidates of *spot* and Wilcoxon ranking are displayed in Figure S4. They reveal strong expression in the *Plasmodium* liver stage as well as weaker expression in trophozoites and oocysts. While the genes performing best in the *spot* ranking show higher expression values in oocysts, the Wilcoxon test ranked genes have more expression in trophozoites. As shown in Figure S2, there is less overall expression in unselected columns of the Wilcoxon ranking, while the *spot* ranked genes have higher expression in the liver stage.

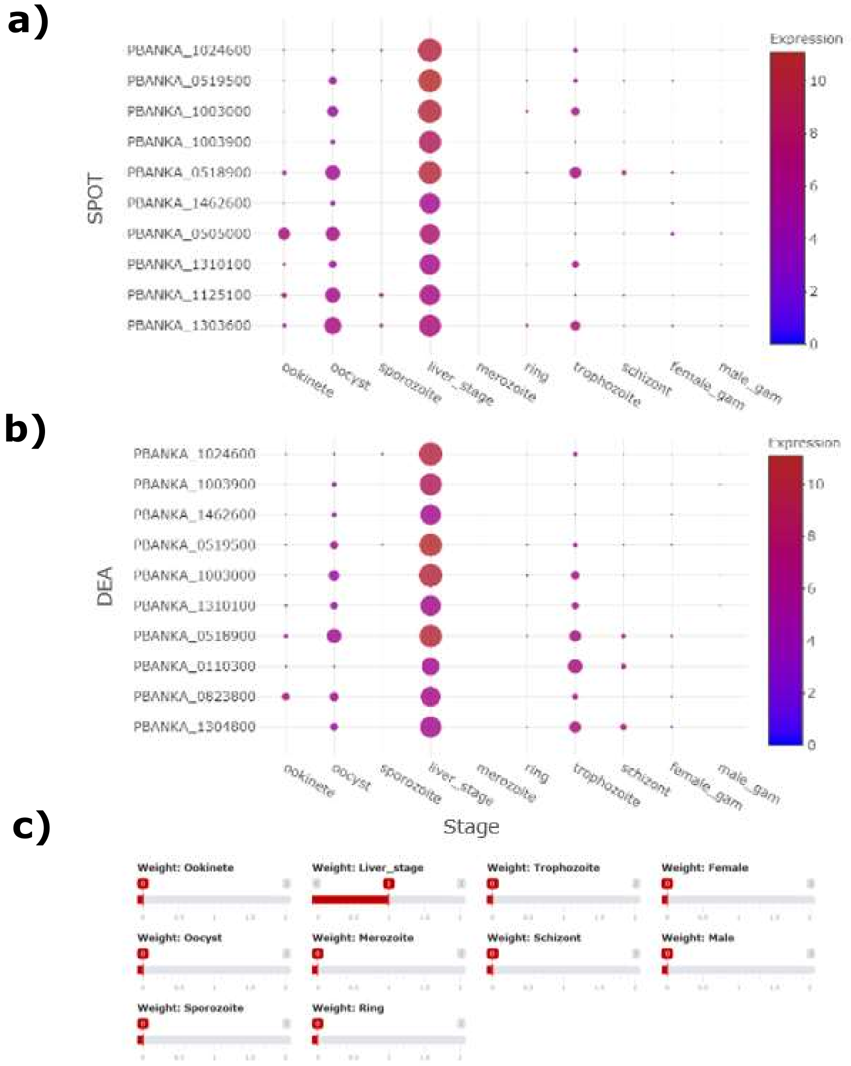

**Figure S4: Comparison of SPOT and DEA reveal higher off-target effects in *spot* ranking**

The 10 best ranked genes from *spot* **(a)** and Wilcoxon ranking **(b).** While the genes of the Wilcoxon ranking have slightly lower values in the unselected columns, *spot* ranked genes have higher values in the liver stage.

### 4 Example prediction B: *Plasmodium* sexual stages

#### 4.1 Purpose

Drug development against malaria focuses mainly on the blood stages of malaria, as these cause the symptoms of the disease and are easily accessible in the laboratory. Since the eradication of malaria has come into the focus of drug design, drugs are also developed against sexual stages to prevent transmission to the mosquito or against liver stages to prevent development of disease-causing blood stages (Kappe, et al., 2010). Drugs targeting the liver stages are routinely used to cure *Plasmodium vivax* infections (Flannery, et al., 2018). However, there are no specific drugs available that target gametocytes. Identifying additional target proteins is therefore of interest (The mal, 2011). Here were present a SPOT search for genes only expressed in sexual blood stages, leaving a high probability for functionality in these stages.

#### 4.2 Results

**Table S3: Results from *spot* ranking for sexual stage specific genes**

| *P. Berghei* GeneID | Gene product description | *spot* score |
|---|---|---|
| PBANKA_1329100 | Plasmepsin VIII | 4,95 |
| PBANKA_0600600 | NIMA related kinase 3, putative | 4,95 |
| PBANKA_1432200 | male development gene 1 | 4,79 |
| PBANKA_1429100 | conserved Plasmodium protein, unknown function | 4,78 |
| PBANKA_1449000 | microgamete surface protein MiGS, putative | 4,49 |
| PBANKA_1361600 | E1-E2 ATPase, putative | 4,25 |
| PBANKA_1038200 | nuclear formin-like protein MISFIT | 4,22 |
| PBANKA_0812600 | conserved Plasmodium protein, unknown function | 4,18 |
| PBANKA_1359600 | 6-cysteine protein | 4,12 |
| PBANKA_1109600 | conserved Plasmodium protein, unknown function | 4,07 |

Sexual stage specific *spot* ranking revealed 3 unknown proteins as well as a wide variety of enzymes, surface proteins and proteins involved in osmiophilic body formation. *P. falciparum* Plasmepsin VIII is known to be active specifically in gametocytes while the p48/45 protein (PBANKA_1359600) is a well characterized surface protein and still in consideration as a target for transmission blocking vaccines (Acquah, et al., 2019; Jiang, et al., 2020; Lee, et al., 2020; van Dijk, et al., 2001; Weißbach, et al., 2017).

Two other interesting candidates are associated with the emergence of osmiophilic bodies, specialized vesicles essential for parasite egress from blood cells and another one was shown to be important in the cell cycle in ookinetes and oocysts (Bushell, et al., 2009; Kehrer, et al., 2016; Ponzi, et al., 2009).

It is therefore hard to predict the function or localization of the 3 unknown genes. However, the interesting phenotypes of proteins with a similar transcription profile suggests that these proteins may have a functional role in mosquito infection as well.

#### 4.3 Comparison to DEA

Similar to the previous section, all genes ranked in the Top 100 of *spot* algorithm are significantly upregulated exclusively in the selected sexual stages (Figure S5 a). In contrast to the expression profiling of liver stages, however, approximately only half of the genes in the top 20 still overlap (Figure S5 b).

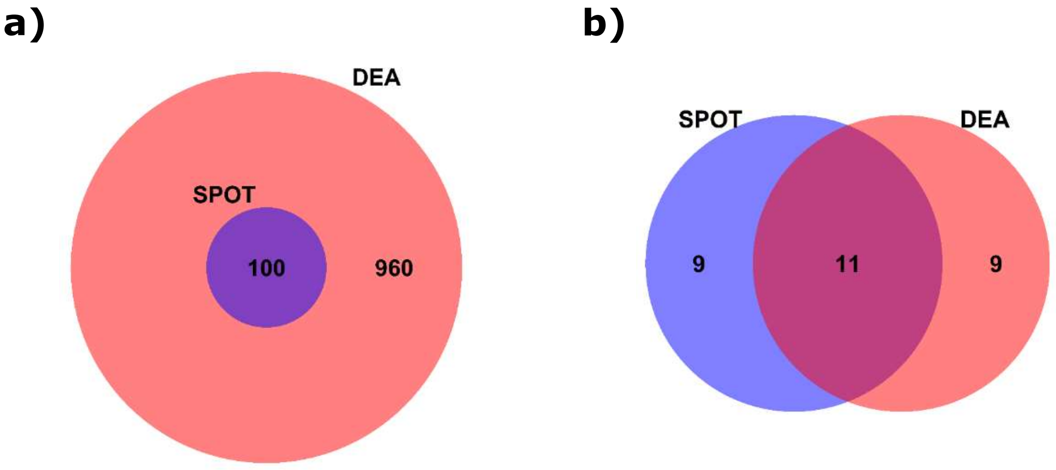

**Figure S5: The majority of top SPOT sex specific genes overlap with DEA results**

**(a)** All genes ranked in the *spot* Top 100 overlap with genes detected as significantly upregulated in the sexual stages (960) **(b)** Comparison of top 20 genes derived from *spot* or the Wilcoxon Rank Sum test (DEA) reveal an overlap of 11 genes.

While the genes in both rankings have very low values in the unselected stages, there are substantially higher expression values in selected stages of the genes ranked by *spot* compared to those ranked by Wilcoxon (Figure S6).

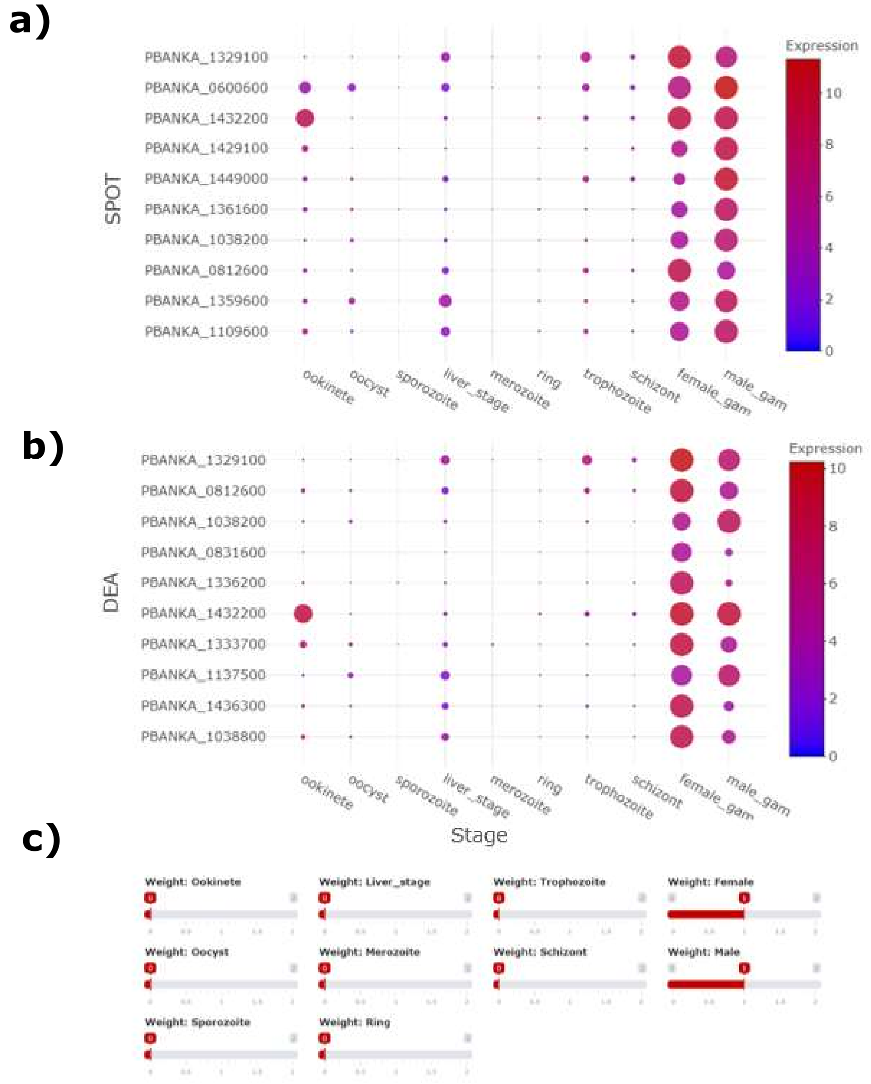

#### Figure S6: *spot* ranked genes reveal high expression in sexual stages

Genes ranking high in the *spot* algorithm **a)** have substantially higher expression values in sexual stages than Wilcoxon ranked genes shown in **b).** Similar to the liver stage results (Figure S4) genes ranking high in the Wilcoxon test have slightly lower expression values in the unselected columns than genes ranked by *spot*.

### 5 Example prediction C: Genes expressed in asexual and sexual blood stages of *Plasmodium*

#### 5.1 Purpose

Asexual and sexual blood stages share the red cell as a host, yet their biology differs dramatically as has also been shown by shifting expression profiles. Here we present a prediction of genes only active in asexual or sexual blood stages but not in other life cycle stages. This should counter select genes involved in general replication mechanisms.

#### 5.2 Results

The results of the third example prediction shown in Table S4 display a multitude of membrane proteins or proteins associated with membrane trafficking and virulence. While the membrane associated histidine-rich protein 1a (MAHRP1a) and the skeleton-binding protein 1 (SBP1) are known to play a role in the transport of parasite proteins to the surface (Blisnick, et al., 2000; De Niz, et al., 2016; Maier, et al., 2007; Spycher, et al., 2003), the function of the ETRAMP protein family is more diverse and needs further evaluation (MacKellar, et al., 2011; Spielmann, et al., 2003). This also applies to the fam-b protein and the p1/s1 nuclease, about which little is known, as well as to the two unknown proteins that are candidates for further analysis.

**Table S4: Results from *spot* ranking of blood and sexual stage specific genes**

| *P. Berghei* GeneID | Gene product description | *spot* score |
|---|---|---|
| PBANKA_1145800 | membrane associated histidine-rich protein 1a | 5,61 |
| PBANKA_1000600 | erythrocyte membrane antigen 1 | 4,96 |
| PBANKA_0300600 | Plasmodium exported protein, unknown function | 4,78 |
| PBANKA_0524800 | early transcribed membrane protein | 4,64 |
| PBANKA_1101300 | skeleton-binding protein 1 | 4,52 |
| PBANKA_0517000 | early transcribed membrane protein | 4,13 |
| PBANKA_0316300 | fam-b protein | 3,62 |
| PBANKA_1200600 | Plasmodium exported protein, unknown function | 3,51 |
| PBANKA_0524200 | early transcribed membrane protein | 3,47 |
| PBANKA_1030600 | p1/s1 nuclease, putative | 3,47 |

#### 5.3 Comparison to DEA

In contrast to the previous predictions, a substantially lower number of genes was significantly upregulated according to the Wilcoxon test (Figure S7 a). Nevertheless, the large majority of genes derived from *spot* ranking still overlapped with these genes. The overlap between the top 20 genes from Wilcoxon and *spot* ranking is 50 %.

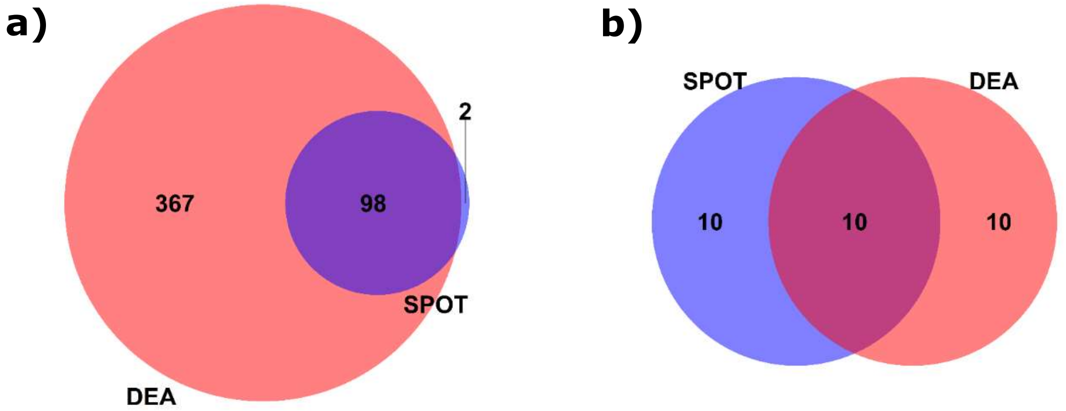

**Figure S7: In more complex searches results from *spot* ranking are still differentially expressed**

**a)** 98 of 100 genes ranked in the *spot* Top 100 overlap with genes detected as significantly upregulated in the sexual stages (367). 367 is by far the lowest number of significantly expressed genes in the example predictions. **b)** 50 % of top 20 genes derived from *spot* ranking or the Wilcoxon Rank Sum test overlap.

However, the genes scoring best in the Wilcoxon test ranking show very high expression in unselected stages compared to all other predictions (Figure S2; Figure S8 b). While the expression values from *spot* ranking derived genes stay low compared to other predictions, they show relatively low expression in the selected stages (Figure S2; Figure S8 a)).

**a)**

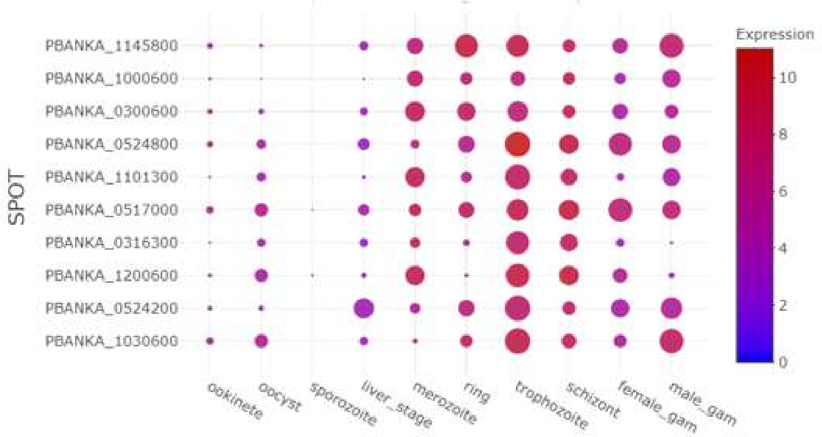

**b)**

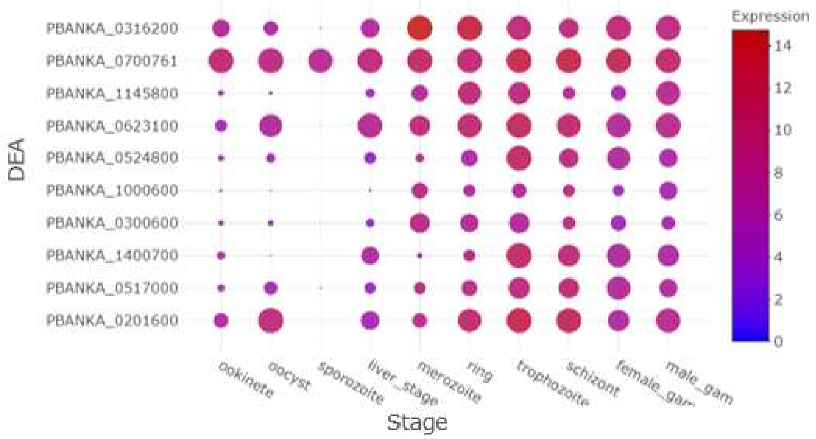

**c)**

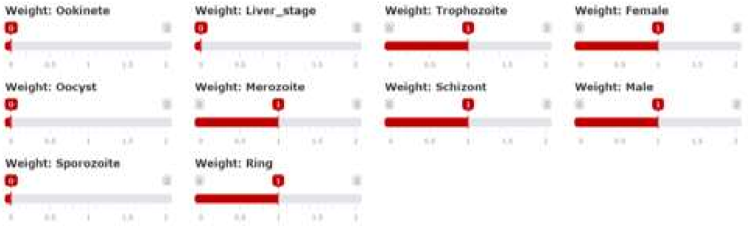

**Figure S8: *spot* ranking for transmission targets shows low expression in unselected stages**

**a)** Genes ranking in the top 10 of *spot* ranking have substantially lower gene expression values in unselected columns compared to genes ranking in the top 10 in the Wilcoxon test **b)**. However, Wilcoxon test ranked genes shown in **b)** have substantially higher expression values in selected stages. Additionally, genes derived by the Wilcoxon test show a higher variance in both categories.

### 6 Requirements

**Table 5: Packages used for web-tool**

| Package | Version |
|---|---|
| DESeq2 | 1.30.0 |
| DT | 0.16 |
| EnvStats | 2.4.0 |
| Limma | 3.46.0 |
| MAST | 1.16.0 |
| matrixStats | 0.57.0 |
| Plotly | 4.9.2.1 |
| Plyr | 1.8.6 |
| Reshape2 | 1.4.4 |
| Rlist | 0.4.6.1 |
| Scales | 1.1.1 |
| Seurat | 3.2.2 |
| Shiny | 1.5.0 |
| Shinybusy | 0.2.2 |
| Shinycssloaders | 1.0.0 |
| shinyEffects | 0.1.0 |
| Shinywidgets | 0.5.4 |
| Sortable | 0.4.4 |
| Stringr | 1.4.0 |
| Tidyr | 1.3.0 |

### 7 Acknowledgements

We thank Omar Harp and Daniel Beiting for critical reading and constructive suggestions on the manuscript as well as Benedikt Broers for discussions.
